# Metabolic response of *Brevibacterium epidermidis* TRM83610 to NaCl stress and ectoine fermentation optimization

**DOI:** 10.1101/2025.09.18.677089

**Authors:** Tangliang Luo, Yafang Zhao, Lijun Wang, Zhanfeng Xia

## Abstract

To elucidate the metabolic response of *Brevibacterium epidermidis* TRM83610 to NaCl stress and facilitate its industrial application, this study employed metabolomics techniques to analyze changes in intracellular metabolites, particularly compatible solutes, under NaCl concentrations of 0, 5%, 10%, and 15%. The ectoine production capacity was further evaluated and optimized using response surface methodology. Results indicated significant metabolic differences among sample groups, with detection of various secondary metabolites associated with antimicrobial activity and plant growth promotion. Six compatible solutes dominated by ectoine were identified. The metabolic response strategies to NaCl stress included osmostress resistance, oxidative stress resistance, and survival competition strategies. Through response surface optimization, ectoine yield reached 440.60 mg/L, representing a 6.22-fold increase over the initial yield of 70.75 mg/L, demonstrating substantial application potential for this strain. This research enriches our understanding of the metabolic profile of *B. epidermidis* TRM83610, preliminarily reveals its metabolic responses to NaCl stress, and provides a foundational basis for its further development and utilization.

**IMPORTANCE:** Our study reports, for the first time, the detection of Nε-Acetyl-L-lysine in *Brevibacterium epidermidis* and the identification of Azetidomonamide A in a microorganism outside of Pseudomonas aeruginosa. It also represents the first elucidation of this strain’s metabolic response to NaCl stress. This research demonstrates the significant application and research value of *B. epidermidis* TRM 83610 and reveals that its strategy for coping with NaCl stress is multifaceted. This includes employing multi-pathway synergy to regulate multiple compatible solutes for osmostress resistance, producing antioxidant compounds for oxidative stress resistance, and secreting antimicrobial compounds as part of a survival competition strategy.

## INTRODUCTION

Under the backdrop of the Fourth Industrial Revolution, traditional chemical engineering-based manufacturing increasingly conflicts with sustainable development principles. In contrast, next-generation industrial biotechnology (NGIB) offers an eco-friendly solution to reduce production costs, conserve energy, mitigate emissions, and enable efficient biosynthesis^[1, 2]^. NGIB employs extremophilic microorganisms as chassis cells for biomanufacturing, with halophilic/halotolerant microbes gaining significant attention in recent years for green production strategies^[2]^.

To date, researchers have developed multiple chassis cells from the genus *Halomonas* for novel biomanufacturing. *Halomonas bluephagenesis* TD01, a promising microbial cell factory, completed pilot-scale trials in 2024 for polyhydroxyalkanoate (PHA) production, demonstrating immense potential for low-cost, large-scale PHA synthesis^[3]^. Ma et al. ^[1]^enhanced ectoine yield to 28 g/L in *Halomonas bluephagenesis* TD-ADEL-58 by integrating chromosomal *ectABC*, *lysC*, and *asd* genes, knocking out degradation-related genes, and implementing dynamic flux regulation via the LuxR-AHL and T7-like orthogonal systems. While the development of halophilic/halotolerant chassis cells underpins NGIB advancement^[4]^, studies on ectoine biosynthesis remain predominantly focused on *Halomonas*, with limited exploration of other genera.

*Brevibacterium epidermidis* has been proposed for amidase production and environmental bioremediation^[5–9]^, highlighting its industrial relevance. However, its metabolic profile—particularly its metabolite responses to NaCl stress and associated regulatory networks—remains poorly characterized.

*B. epidermidis* TRM83610, a halotolerant strain isolated from Mangya Emerald Lake on the Qinghai-Tibet Plateau, exhibits stable cellular morphology at 0–15% NaCl^[10]^, robust growth at 30–40°C^[11]^. This adaptability to environmental fluctuations may have driven the evolution of unique metabolic mechanisms, enabling the strain to produce industrially valuable metabolites, demonstrate a robust capacity for compatible solute accumulation^[12, 13]^, and potentially synthesize diverse compatible solutes^[14–16]^.

In this study, we employed metabolomics to analyze intracellular metabolite levels in *B. epidermidis* TRM83610, under varying NaCl concentrations, with particular focus on abundance changes of compatible solutes supplemented by targeted metabolomic analysis. Additionally, fermentation conditions were optimized using response surface methodology to enhance ectoine production yield. This approach facilitates the elucidation of the strain’s metabolic regulatory network, thereby advancing understanding of its metabolic mechanisms for salt stress tolerance and promoting industrial applications. The research aims to reveal the metabolic response of *B. epidermidis* TRM83610 to NaCl stress while evaluating its ectoine production capacity, thereby establishing a foundational basis for future metabolic engineering efforts to develop this strain into a chassis cell for industrial utilization.

## RESULTS

### Non-targeted metabolomics analysis

#### Quality control evaluation

As shown in Fig. 1, the tight clustering of QC samples with minimal inter-sample variation confirmed the stability of the analytical methodology and instrumentation, ensuring reliable metabolite detection. Within-group samples for both control and treatment groups clustered within the 95% confidence interval, exhibiting low intra-group variability, while distinct inter-group separation underscored statistically significant metabolic differences between conditions.

**Fig 1.**
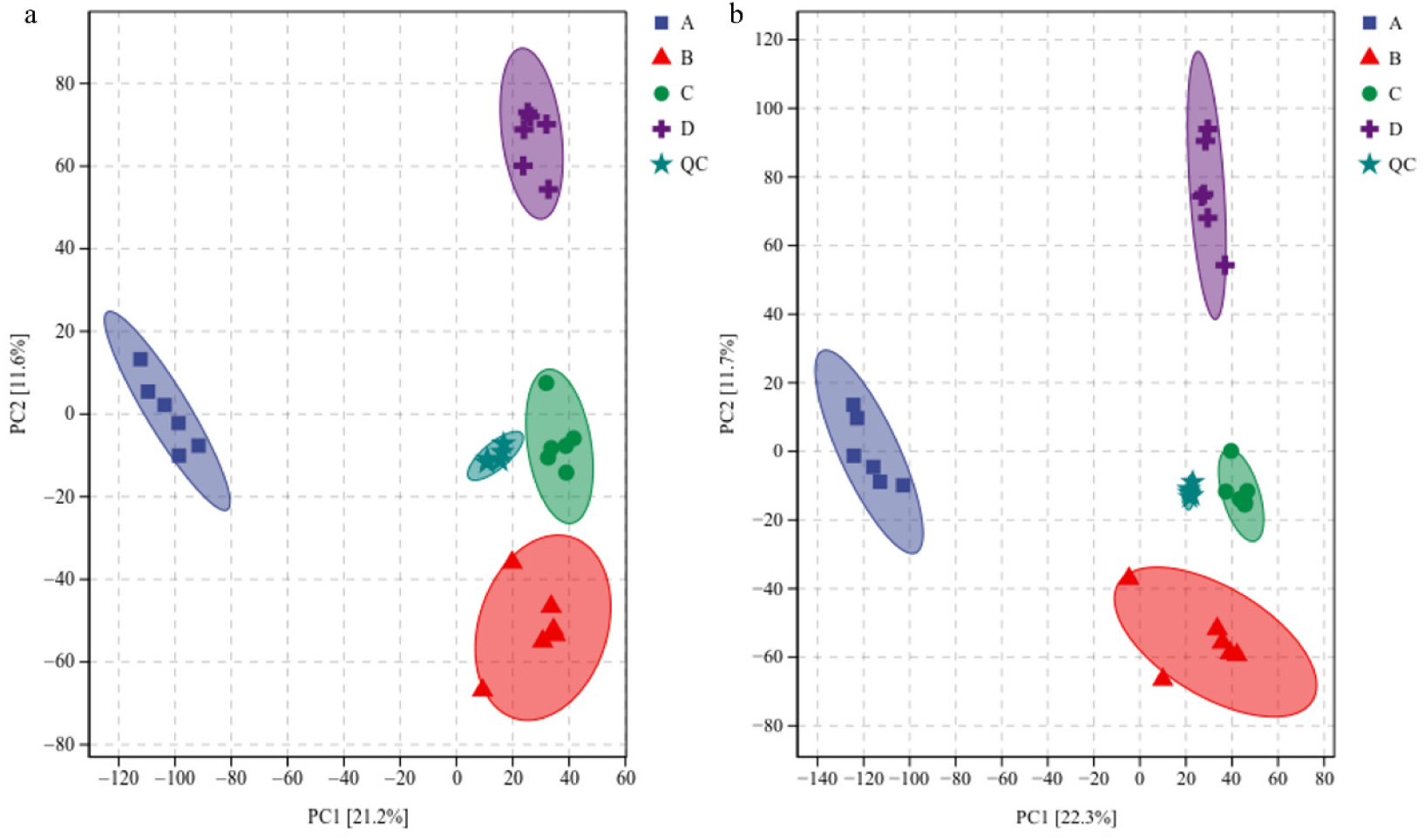
Principal component analysis (PCA) overview. (a) PCA in positive ion mode; (b) PCA in negative ion mode. Labels A-D denote NaCl concentrations of 0, 5%, 10%, and 15%, respectively.

#### Impact of NaCl Stress on Intracellular Metabolites in *B. epidermidis*

Metabolite identification via database matching using retention time, mass-to-charge ratio (m/z), and molecular weight revealed 1,416 metabolites, including 985 in positive ion mode and 431 in negative ion mode. Metabolites are primarily classified into nine major categories (Fig. 2a, b). In positive ion mode (Figure 2a), the predominant metabolites were organic heterocyclic compounds (29.5%), organic acids and derivatives (21.9%), benzenoids (15.3%), lipids and lipid-like molecules (11.4%), oxygen-containing organic compounds (6.1%), phenylpropanoids and polyketides (5.7%), and nitrogen-containing organic compounds (5.5%). In negative ion mode (Figure 2b), the predominant metabolites were organic acids and derivatives (25.2%), lipids and lipid-like molecules (22.1%), organic heterocyclic compounds (19.3%), benzenoids (15.0%), phenylpropanoids and polyketides (6.5%), and oxygen-containing organic compounds (5.4%).

**Fig 2.**
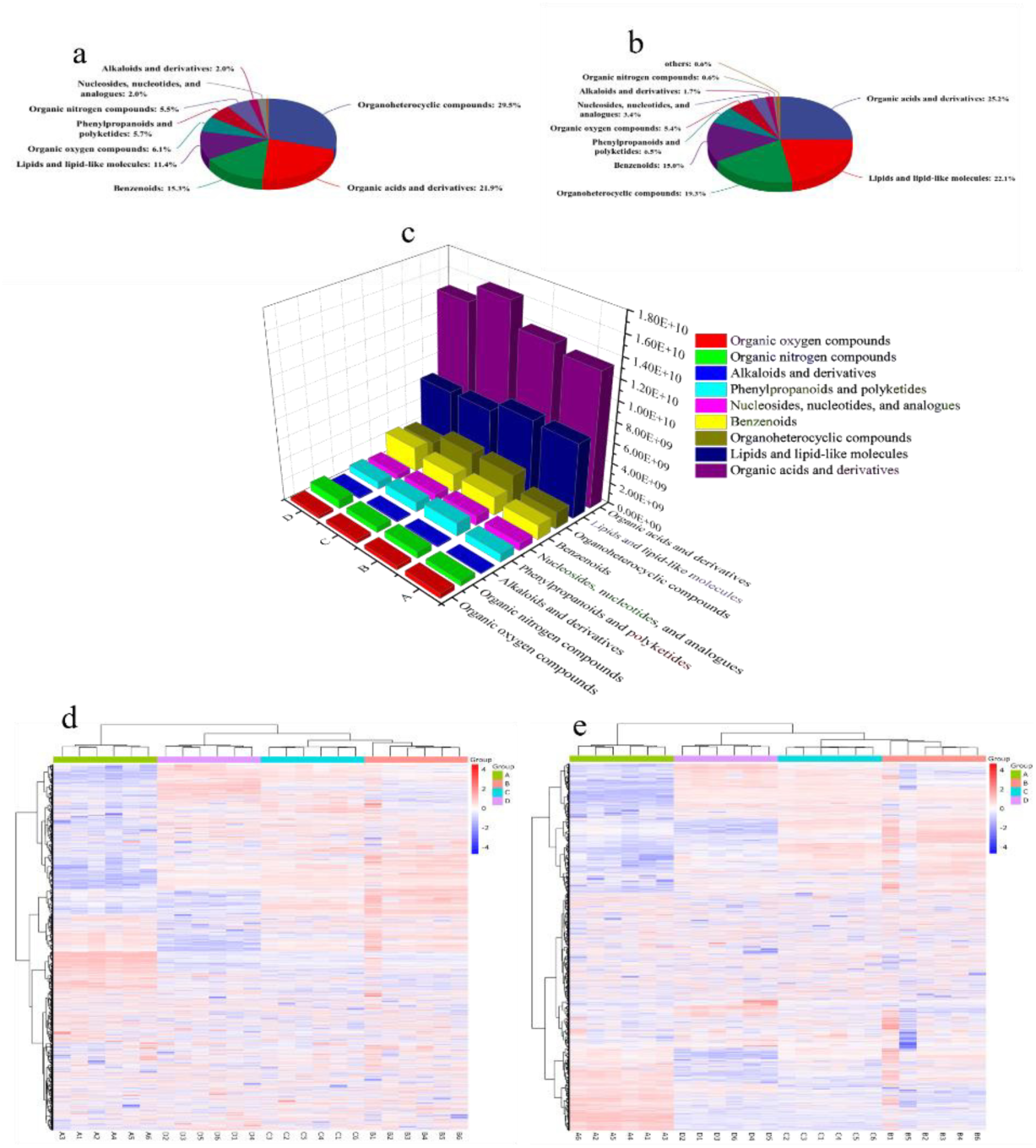
Differential metabolite distribution and hierarchical clustermg analysis. (a) Superclass distribution of DMs in positive ion mode; (b) superclass distribution of DMs in negative ion mode; (c): histogram of relative abundance changes for 9 compound classes; (d) hierarchical clustering in positive ion mode; (e) hierarchical clustering in negative ion mode. LabelsA-D correspond to NaCl concentrations of 0, 5%, 10%, and 15%.

NaCl stress significantly altered the metabolic profile of *B. epidermidis* TRM83610. The total relative abundance of nine major metabolite classes was affected by NaCl stress (Fig. 2c). When the NaCl concentration increased, the abundance of Organic oxygen compounds gradually decreased, while the abundance of Benzenoids gradually increased. The abundances of the remaining classes all showed an initial increase followed by a decrease. Significant changes occurred in the relative abundance of intracellular metabolites across different NaCl concentrations (Fig. 2d, e). Hierarchical clustering heatmap analysis revealed that the sample groups formed two distinct clusters: an initial branch without NaCl addition, and a second branch containing samples exposed to 5%, 10%, and 15% NaCl. Notably, the samples treated with 5% and 10% NaCl clustered closely together, indicating a high degree of similarity in their metabolic profiles, with significant but relatively small differences between them. This suggests that the metabolic response of the strain to 5% and 10% NaCl stress was similar, as evidenced by comparable numbers of differential metabolites (DMs). However, the relative abundances of metabolites still changed progressively with increasing NaCl concentration.

#### Pairwise Comparative Differential Analysis

Orthogonal partial least squares-discriminant analysis (OPLS-DA) permutation test plots (Fig. S1) demonstrated that all Q² values for permuted models fell below the original Q² value (far right), confirming the absence of overfitting and validating the model’s reliability. DMs analysis were subsequently performed using this robust model. DMs were filtered based on *P* < 0.05 and fold change >2 or <0.5, then visualized via multi-group differential volcano plots (Fig. 3). Upregulated DMs progressively increased from left to right (Table 1). Total DMs initially rose and then declined with escalating NaCl concentrations, peaking at 10% NaCl (642 DMs) and 15% NaCl (632 DMs), with 434 and 422 DMs in positive ion mode and 208 and 210 in negative ion mode, respectively. The 5% vs. 10% NaCl comparison yielded the fewest DMs (189 total: 141 in positive, 48 in negative), indicating minimal metabolic divergence between these groups. Conversely, the control (0 NaCl) vs. 10% NaCl comparison exhibited the highest number of DMs, underscoring the strain’s strongest metabolic response under 10% NaCl stress.

**Fig 3.**
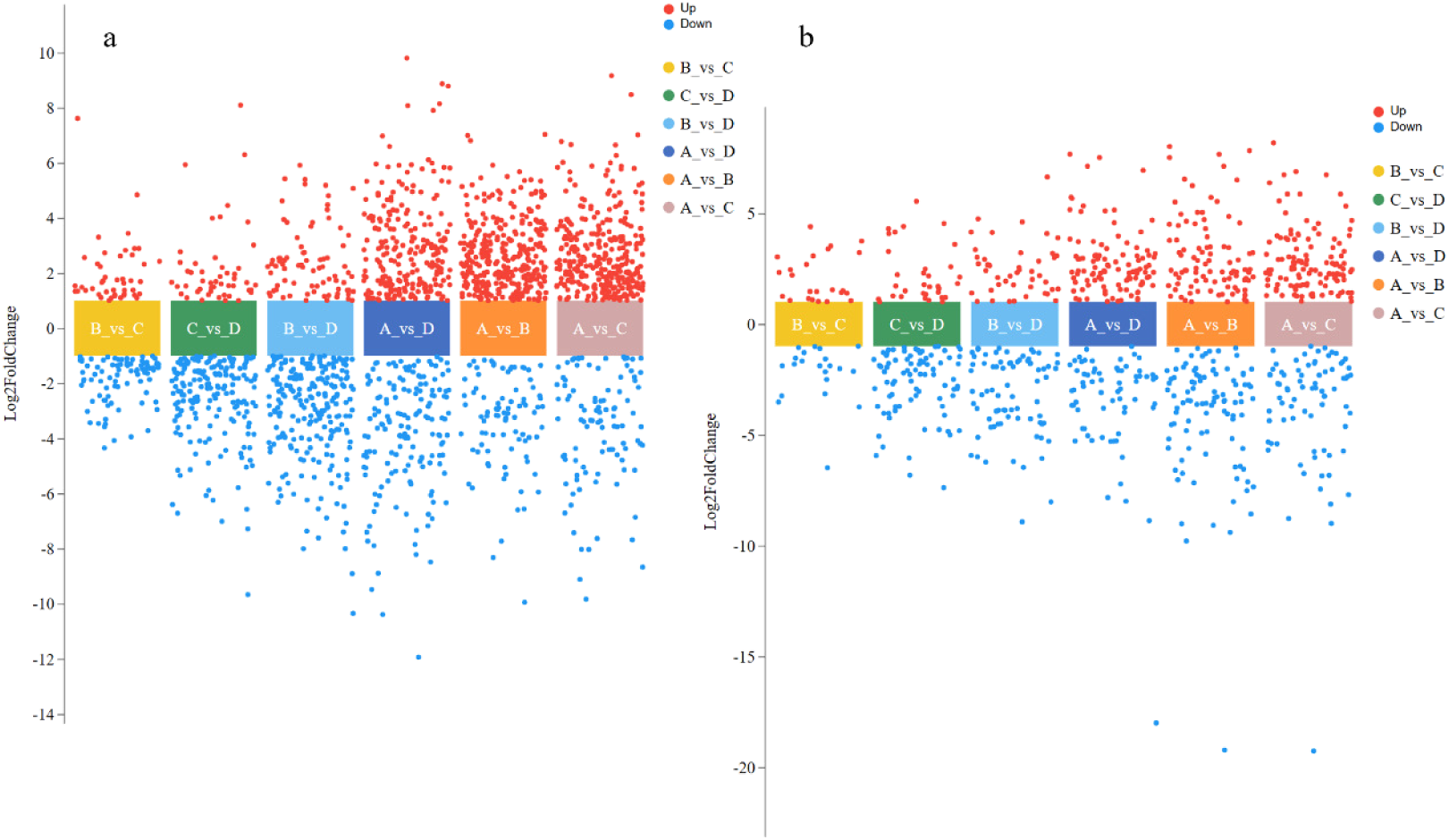
Multi-group differential volcano plots. Each dot represents a DM, with red indicating upregulation and blue downregulation. (a) Positive ion mode; (b) Negative ion mode. Labels A-D: NaCl concentrations of 0, 5%, 10%, and 15%. Comparisons (e.g., A vs B) denote pairwise analysis between groups Band A.

**Table 1.**
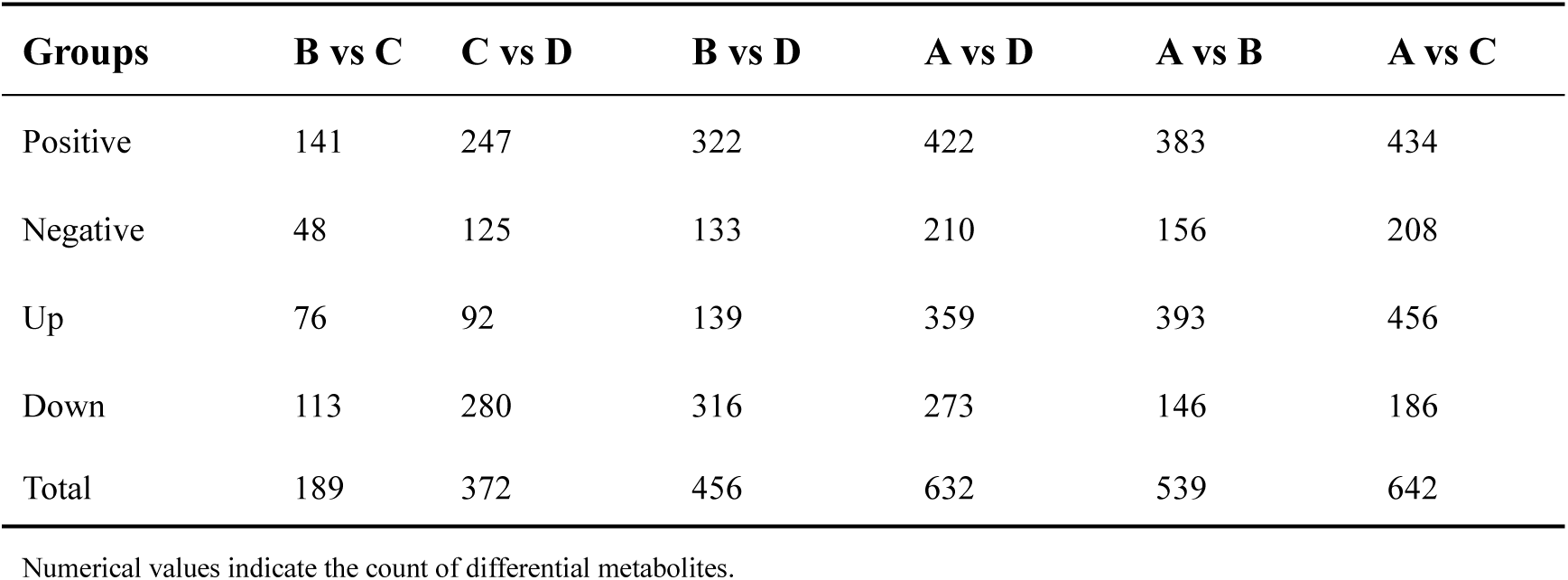
Number of DMs across sample groups.

According to the criteria of *P* < 0.01, FDR < 0.05, fold change > 2 or < 0.5, and VIP > 1, differentially abundant metabolites (DMs) were screened. The Venn diagram of DMs (Fig. 4) revealed that in positive ion mode, the treatment groups shared 182 DMs, with 82 DMs common between 5% NaCl and 10% NaCl, 22 DMs common between 5% NaCl and 15% NaCl, and 52 DMs common between 10% NaCl and 15% NaCl. In negative ion mode, the treatment groups shared 98 DMs, with 21 DMs common between 5% NaCl and 10% NaCl, 15 DMs common between 5% NaCl and 15% NaCl, and 31 DMs common between 10% NaCl and 15% NaCl. Analysis of unique DMs in each treatment group showed that in positive ion mode, 10% NaCl had the fewest unique DMs (29), while in negative ion mode, 5% NaCl had the fewest unique DMs (17), followed closely by 10% NaCl with 18 unique DMs.

**Fig 4.**
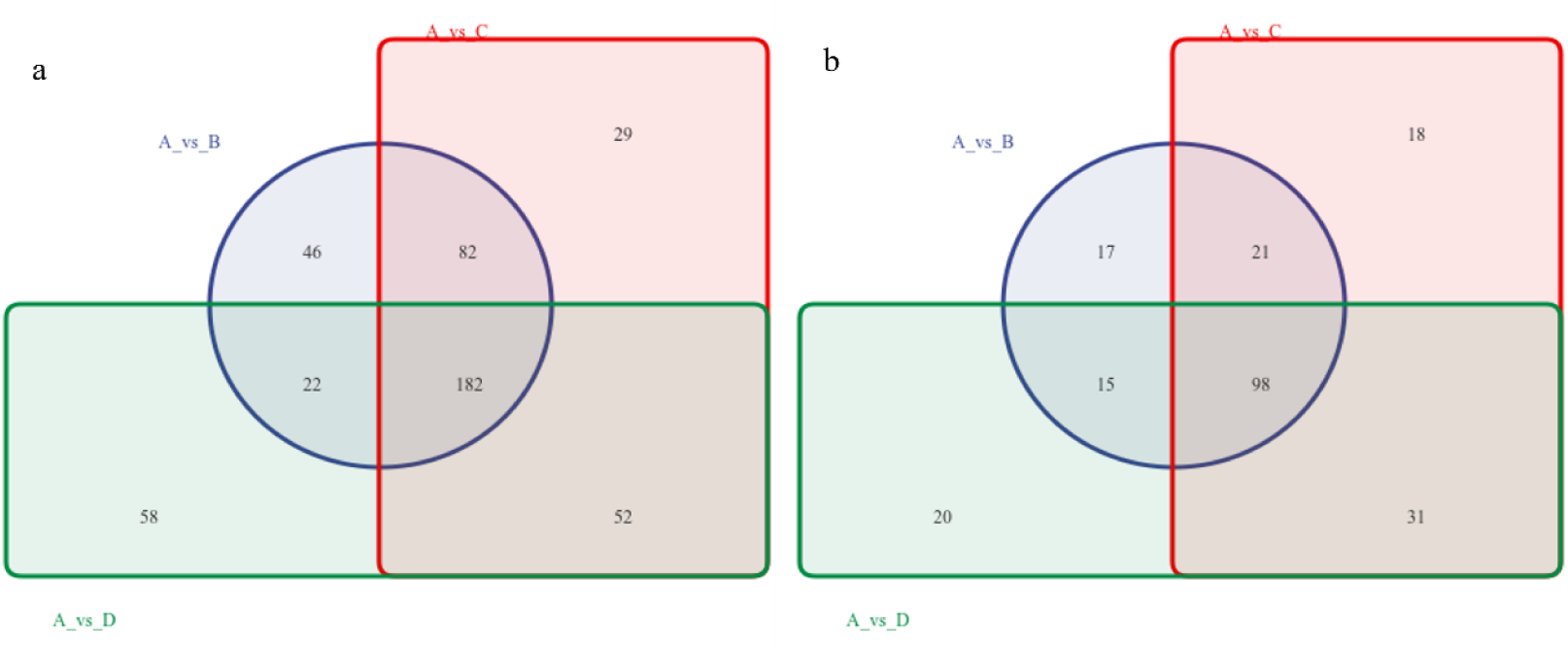
Gear venn diagrams of differential metabolites. (a) Gear Venn diagram in positive ion mode; (b) Gear Venn diagram in negative ion mode. Labels A-D: NaCl concentrations of 0, 5%, 10%, and 15%. Comparisons (e.g., A vs 8) denote pairwise group analysis.

#### KEGG Enrichment Analysis

In the permutation test plot of the partial least squares-discriminant analysis (PLS-DA) (Fig. S2), all Q² points in both positive and negative ion modes were lower than the original Q² point on the far right, indicating that the model was reliable and effective without overfitting. Based on this model, multi-group comparative differential analysis of metabolites was conducted. Given the large number of differentially abundant metabolites (DMs) identified in pairwise comparisons, these abundance changes might not entirely reflect NaCl concentration-responsive DMs, potentially interfering with the analysis process and experimental results and complicating the screening of critical metabolites. Therefore, in the multi-group comparative analysis, KEGG enrichment analysis was first applied to the differential metabolite sets to filter out most metabolites via thresholding.

By leveraging KEGG enrichment analysis, differentially abundant metabolites were mapped to specific metabolic pathways to identify key pathways and analyze their roles under NaCl stress. Subsequently, 22 metabolic pathways were screened from 96 candidate pathways using a threshold of *P* < 0.01 and visualized in a factor loading plot (Fig. 5). As shown in Fig. 5, the most significantly enriched pathway was ABC transporters, followed by Biosynthesis of amino acids. Enriched pathways related to amino acid metabolism included Lysine degradation, Alanine, aspartate and glutamate metabolism, Glycine, serine and threonine metabolism, Arginine biosynthesis, Phenylalanine, tyrosine and tryptophan biosynthesis, and D-Amino acid metabolism. These pathways regulate the synthesis or degradation of specific amino acids, meeting cellular demands for acidic substances under salt stress ^[15]^.

**Fig 5.**
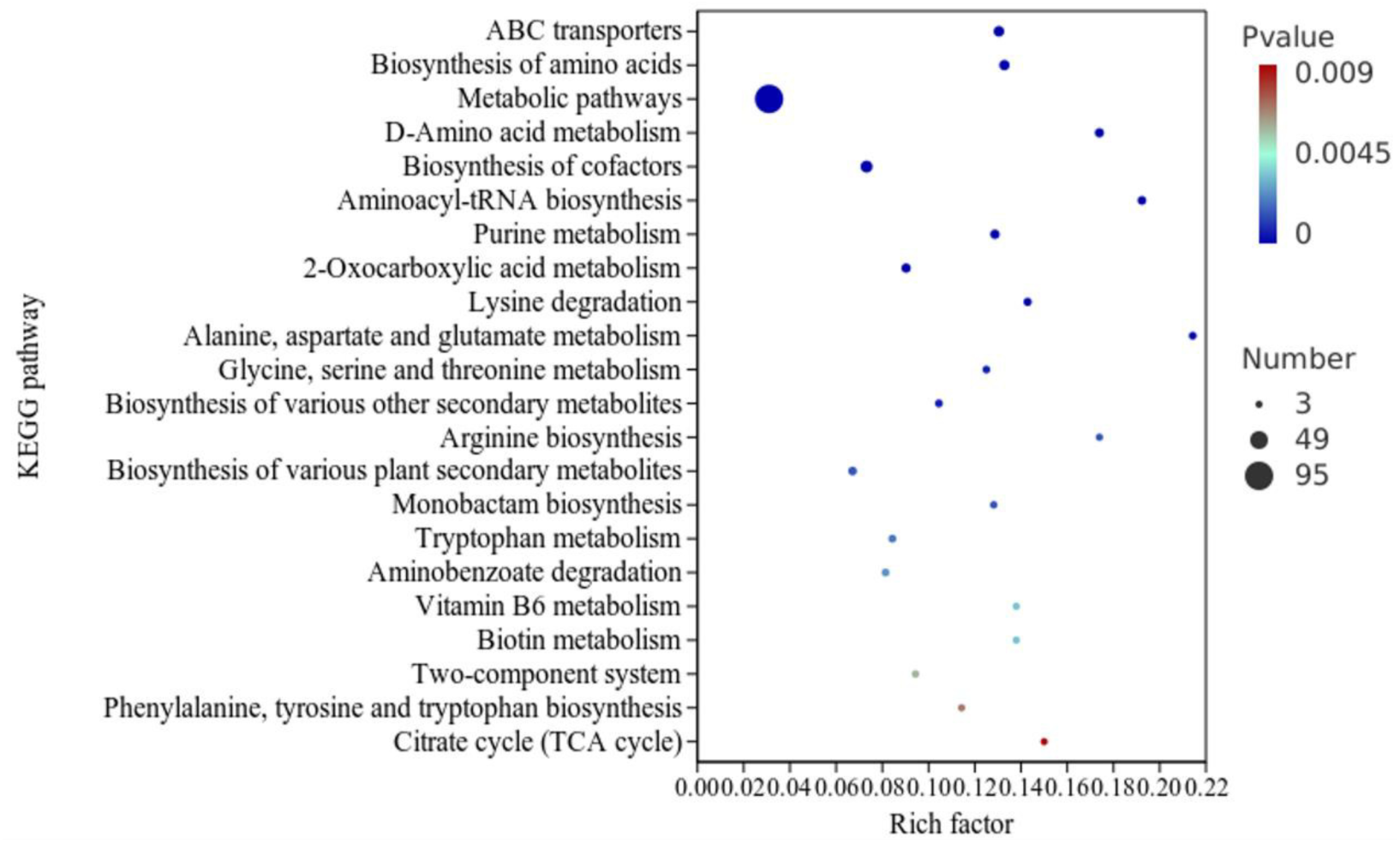
KEGG enrichment analysis bubble plot. X-axis: Enrichment ratio; Y-axis: Metabolic pathways. Color intensity reflects significance (blue: lower; red: higher). Dot size correlates with the number of enriched compounds.

#### Significantly Enriched DMs and Correlation Analysis

A total of 102 DMs were mapped to 22 significantly enriched metabolic pathways (Table S1), encompassing compounds associated with osmoprotection, antioxidant activity, antimicrobial effects, anti-inflammatory properties, and plant growth promotion, highlighting the broad application potential of *B. epidermidis* TRM83610. The relative abundances of these compounds under different NaCl concentrations are presented in Table 2.

**Table 2.**
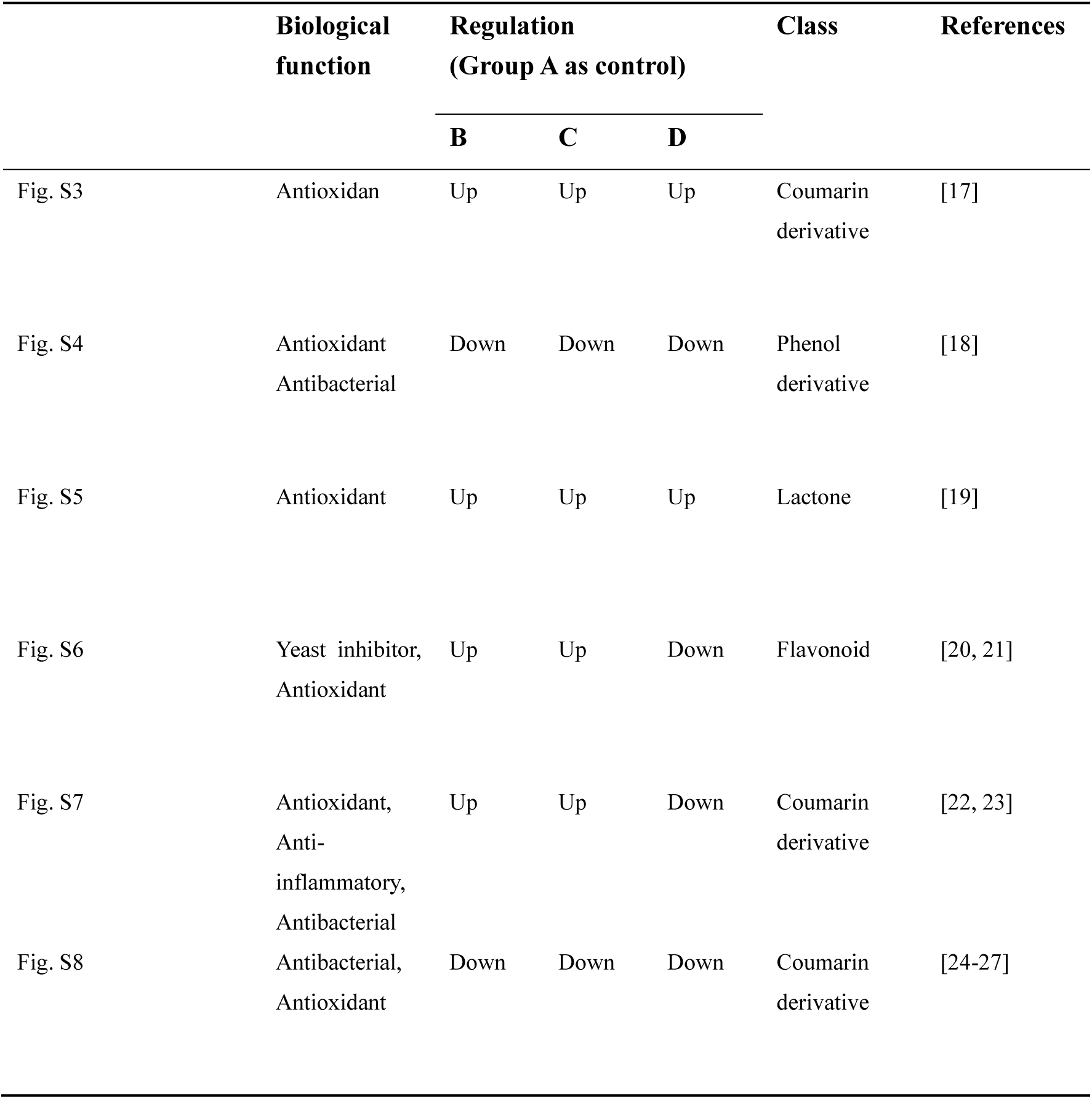
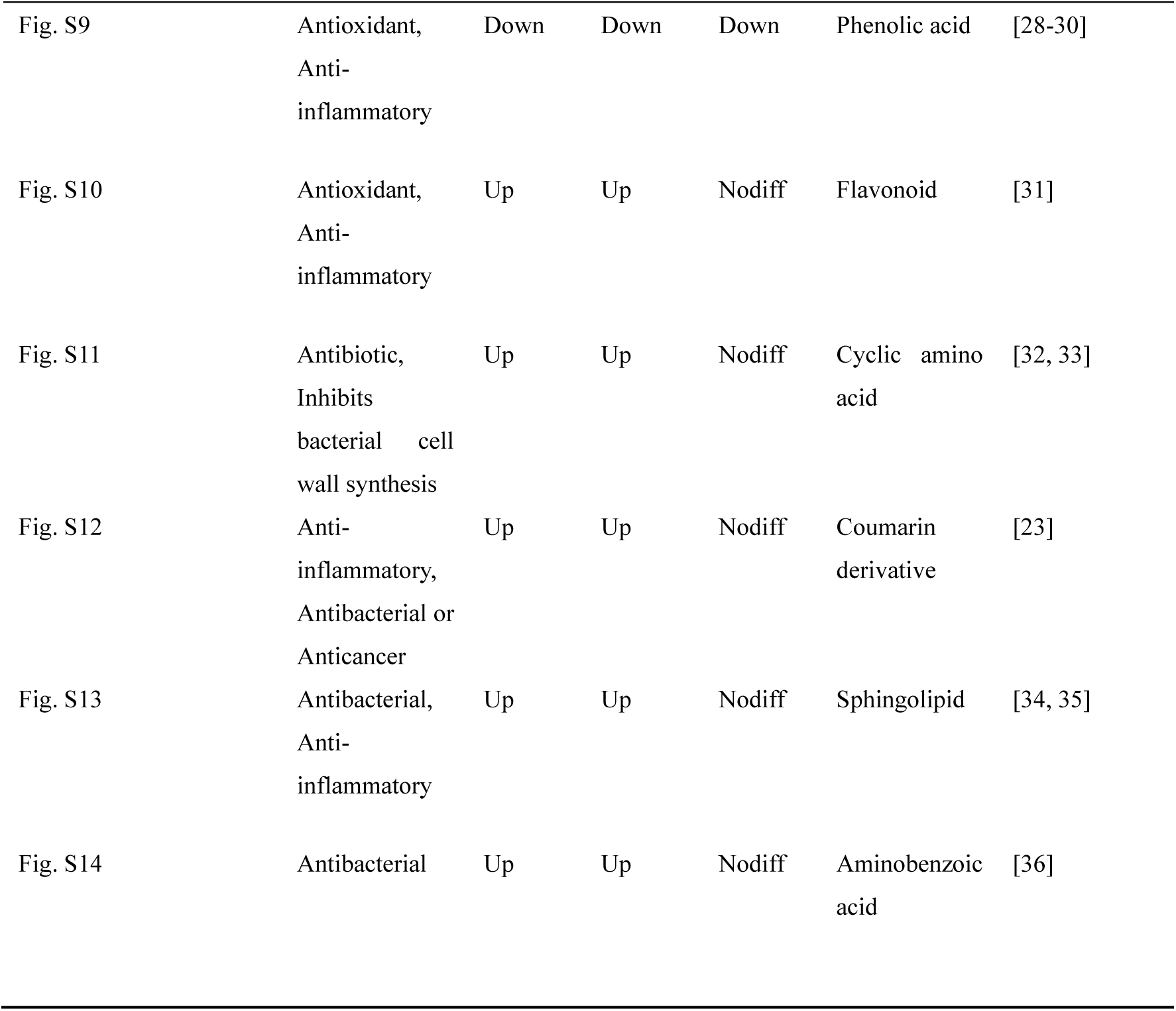
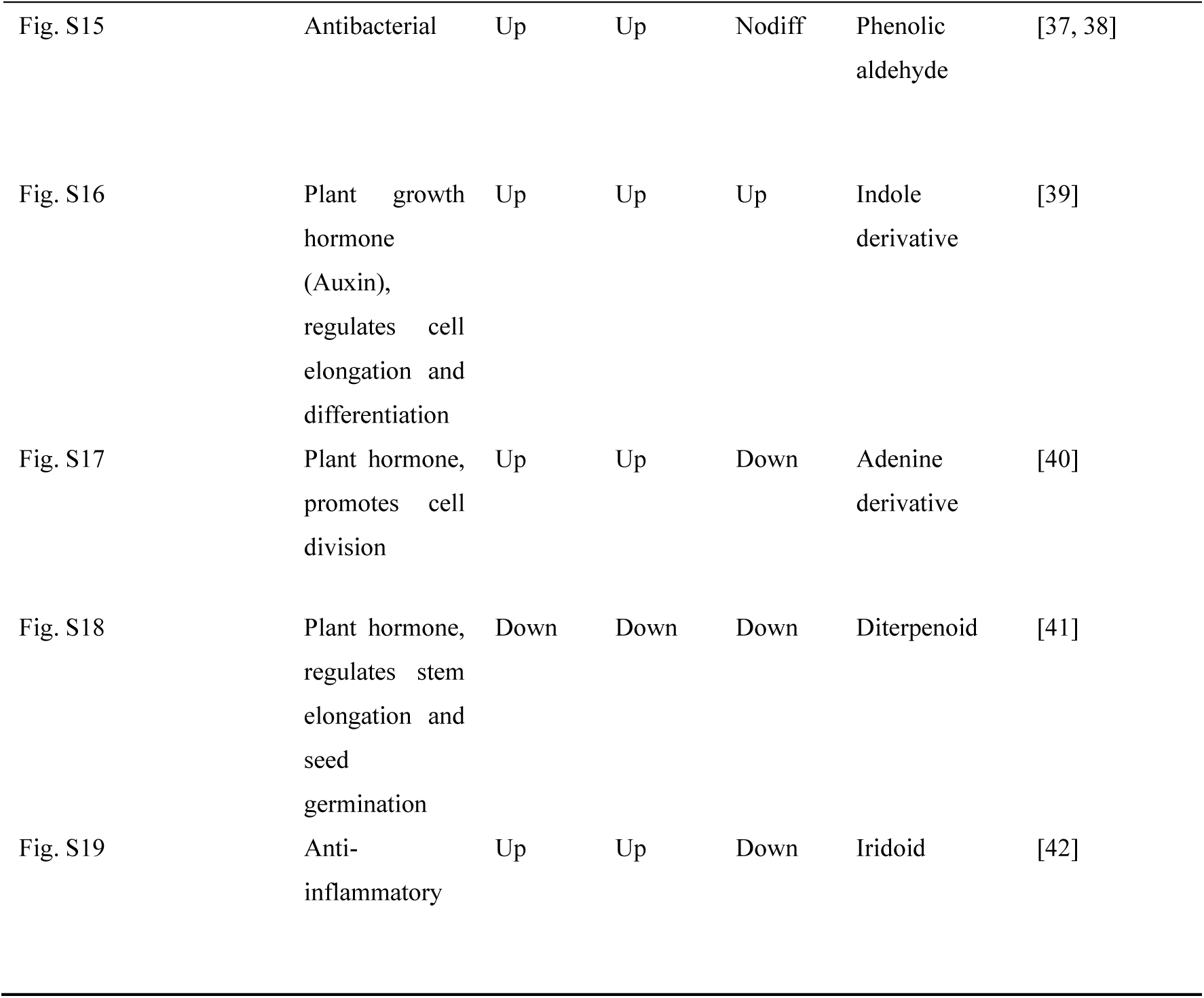
Significantly enriched functional DMs.

|  | Biological<br>function | Regulation<br>(Group A as control) |  |  | Class | References |
| --- | --- | --- | --- | --- | --- | --- |
|  |  | B | C | D |  |  |
| Fig. S3 | Antioxidan | Up | Up | Up | Coumarin<br>derivative | [17] |
| Fig. S4 | Antioxidant<br>Antibacterial | Down | Down | Down | Phenol<br>derivative | [18] |
| Fig. S5 | Antioxidant | Up | Up | Up | Lactone | [19] |
| Fig. S6 | Yeast inhibitor,<br>Antioxidant | Up | Up | Down | Flavonoid | [20, 21] |
| Fig. S7 | Antioxidant,<br>Anti-inflammatory,<br>Antibacterial | Up | Up | Down | Coumarin<br>derivative | [22, 23] |
| Fig. S8 | Antibacterial,<br>Antioxidant | Down | Down | Down | Coumarin<br>derivative | [24-27] |
|  |  | B | C | D |  |  |
| Fig. S9 | Antioxidant,<br>Anti-inflammatory | Down | Down | Down | Phenolic acid | [28-30] |
| Fig. S10 | Antioxidant,<br>Anti-inflammatory | Up | Up | Nodiff | Flavonoid | [31] |
| Fig. S11 | Antibiotic,<br>Inhibits<br>bacterial cell<br>wall synthesis | Up | Up | Nodiff | Cyclic amino<br>acid | [32, 33] |
| Fig. S12 | Anti-inflammatory,<br>Antibacterial or<br>Anticancer | Up | Up | Nodiff | Coumarin<br>derivative | [23] |
| Fig. S13 | Antibacterial,<br>Anti-inflammatory | Up | Up | Nodiff | Sphingolipid | [34, 35] |
| Fig. S14 | Antibacterial | Up | Up | Nodiff | Aminobenzoic<br>acid | [36] |
|  |  | B | C | D |  |  |
| Fig. S15 | Antibacterial | Up | Up | Nodiff | Phenolic<br>aldehyde | [37, 38] |
| Fig. S16 | Plant growth hormone (Auxin), regulates cell elongation and differentiation | Up | Up | Up | Indole<br>derivative | [39] |
| Fig. S17 | Plant hormone, promotes cell division | Up | Up | Down | Adenine<br>derivative | [40] |
| Fig. S18 | Plant hormone, regulates stem elongation and seed germination | Down | Down | Down | Diterpenoid | [41] |
| Fig. S19 | Anti-inflammatory | Up | Up | Down | Iridoid | [42] |

Compatible solutes are pivotal for halotolerant microorganisms to mitigate NaCl stress. Among the significantly enriched differentially abundant metabolites (DMs), six potential compatible solutes were identified: ectoine, betaine, L-glutamic acid, L-glutamine, Nε-acetyl-L-lysine, and L-proline, with ectoine exhibiting a maximum relative abundance significantly higher than the others. These solutes have been validated to function as osmoprotectants in halotolerant or halophilic microbial cells ^[43, 44]^, yet their relative abundance trends under varying NaCl concentrations were inconsistent (Fig. 6). Specifically, as NaCl concentration increased, the relative abundance of ectoine progressively rose, peaking at 15% NaCl. In contrast, Nε-acetyl-L-lysine initially increased before declining, while betaine, L-glutamic acid, L-glutamine, and L-proline showed gradual reductions in relative abundance with elevated NaCl levels.

**Fig 6.**
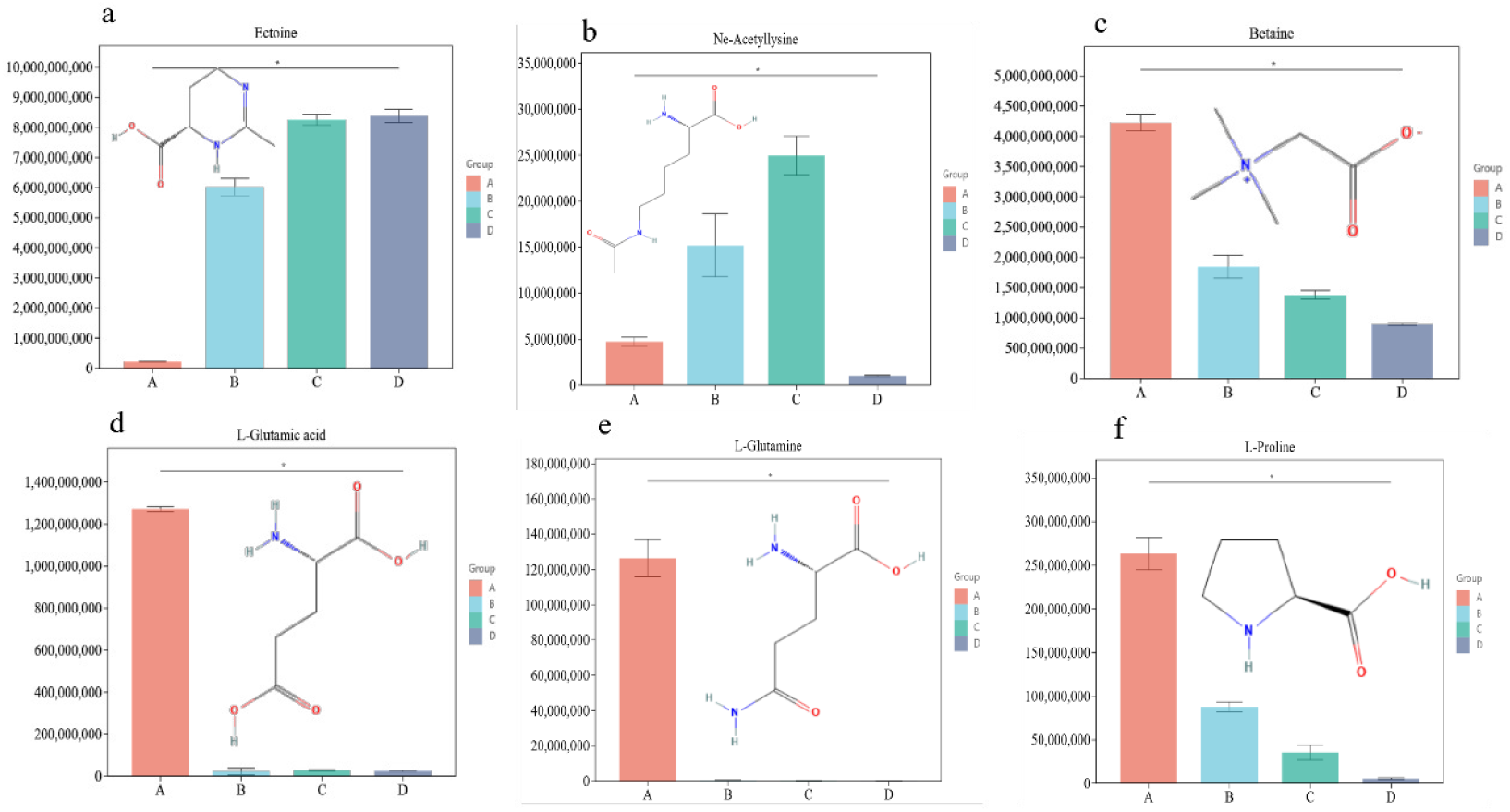
Potential compatible solutes and their abundance changes. Note: (a-f) Bar charts represent the relative abundances of Ectoine, Ni:-acetyl-L-lysine, L-glutamic acid, L-glutamine, L-proline, and betaine, respectively.

To analyze the correlations between significantly enriched differentially abundant metabolites (DMs) and ectoine, thereby revealing the major metabolite classes influencing ectoine synthesis and their associations with other compatible solutes, Spearman correlation coefficient analysis was performed on DMs from the top 22 significantly enriched metabolic pathways (Table S2). The results were visualized through a global correlation network (Fig 7a) and a local correlation network (Fig 7b). Spearman correlation analysis revealed that 31 metabolites exhibited strong negative correlations with ectoine (*r*s< −0.7), including 14 organic acids and derivatives, and 6 nucleosides, nucleotides, and analogs. Conversely, 6 metabolites showed strong positive correlations with ectoine (*r*s > 0.7), among which 3 were indole and its derivatives. As illustrated in Fig. 7a, ectoine and Nε-acetyl-L-lysine displayed negative correlations with betaine, L-proline, L-glutamic acid, L-glutamine, and L-aspartic acid. In contrast, betaine, L-proline, L-glutamic acid, L-glutamine, and L-aspartic acid were positively correlated with each other. Notably, Nε-acetyl-L-lysine showed no significant correlations with other compatible solutes.

**Fig 7.**
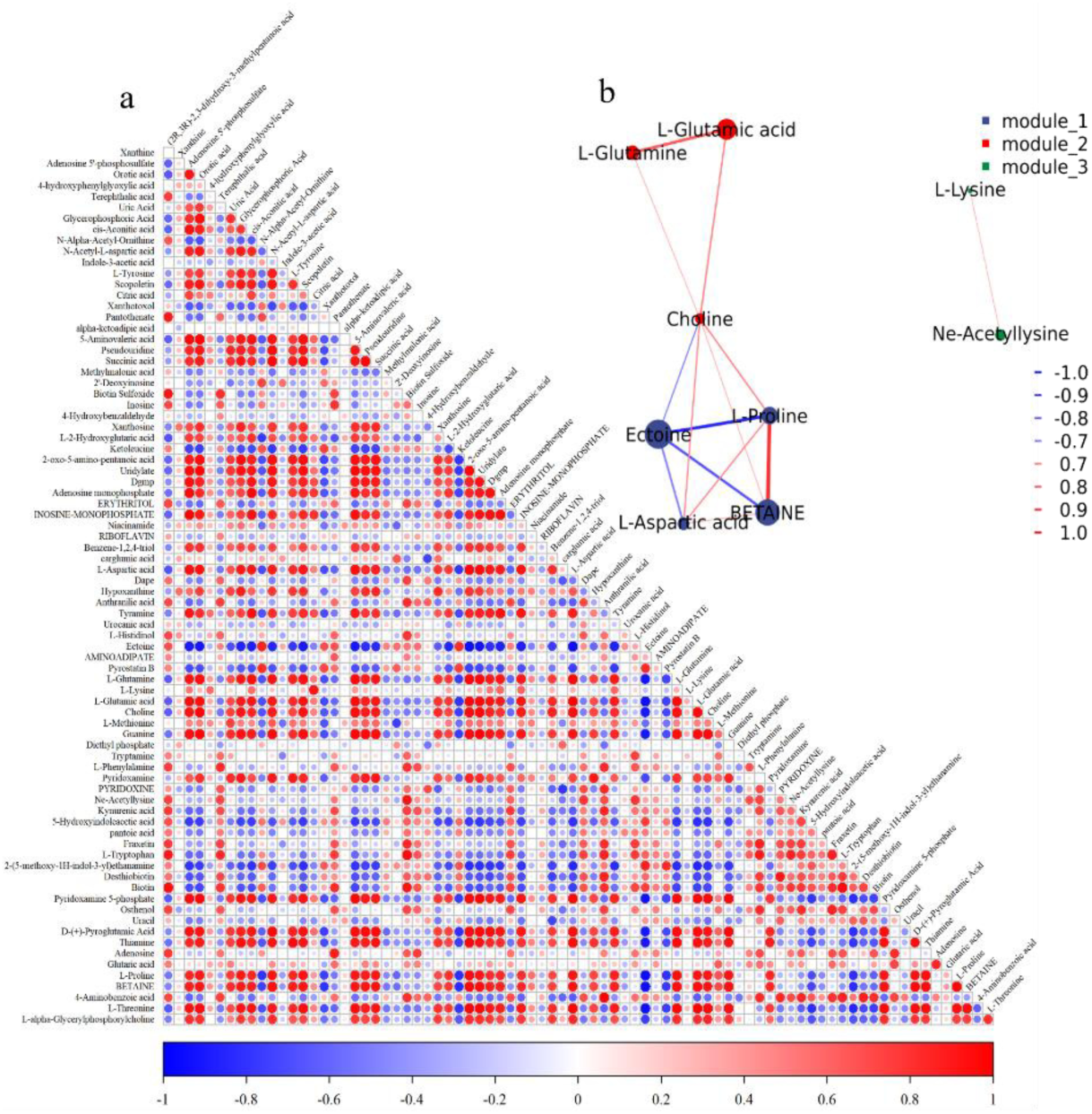
Correlation analysis of differentially abundant metabolites. Red indicates positive correlations, blue indicates negative correlations. (a) Global correlation analysis: darker colors denote stronger correlations. (b) Local correlation network analysis: thicker lines and darker colors represent stronger correlations; node size is proportional to the maximum relative abundance of compounds across different NaCl concentrations.

#### Metabolic Network Analysis

Based on the KEGG enrichment analysis results, a metabolic network diagram of specific metabolites was constructed using metabolite-pathway relationships from KEGG PATHWAY (Fig. 8). The identified compatible solutes were tightly interconnected through multiple metabolic pathways. Cells uptake amino acids from the extracellular environment via ABC transporters. Aspartate serves as th e substrate for ectoine synthesis, while glutamate provides amino groups for ectoine biosynthesis and can also be converted to aspartate as a supplementary source. Glutamine and proline are metabolized into glutamate to replenish its pool. Lysine metabolism generates Nε-acetyl-L-lysine, which undergoes deacetylation to regenerate lysine. Oxidative degradation of lysine ultimately produces succinate, which enters the tricarboxylic acid (TCA) cycle. Choline acts as the substrate for betaine synthesis, and betaine may be converted to glycine as a carbon source, subsequently forming tryptophan ^[45]^, which is furth ermetabolized into 5-hydroxyindoleacetate and 5-methoxytryptamine. Notably, citrate, succinate, and cis-aconitate were significantly downregulated, suggesting an accelerated TCA cycle to provide additional energy for cells under NaCl stress.

**Fig 8.**
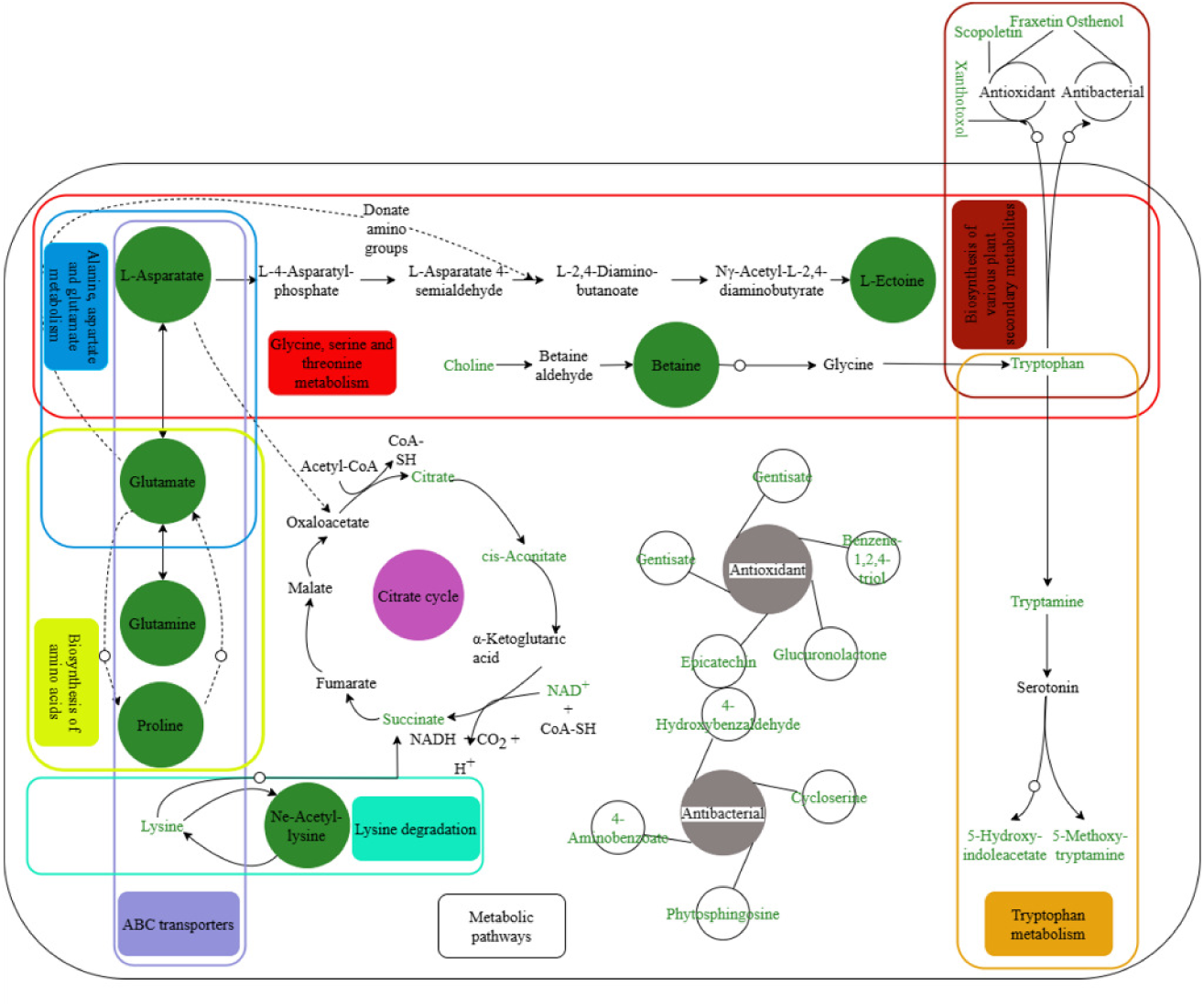
Metabolic network diagram between specific metabolites and KEGG Pathways. Green circles: represent specific metabolites; green text: detected metabolites; solid arrows: reactions defined in KEGG pathways; dashed arrows: putative reactions; circular arrows: reactions involving intermediate products; rounded rectangles with distinct colors denote different metabolic pathways

### Targeted Identification Analysis

#### Targeted identification of intracellular compatible solutes in *B. epidermidis* TRM 83610

Untargeted metabolomics analysis preliminarily indicated that the primary compatible solute in TRM83610 might be ectoine. To confirm this identification, LC-MS was employed to analyze the intracellular extract of TRM83610. Comparative assessment of retention times and mass-to-charge ratios (m/z) with an ectoine standard (Fig. 9) demonstrated that TRM83610 produces ectoine.

**Fig 9.**
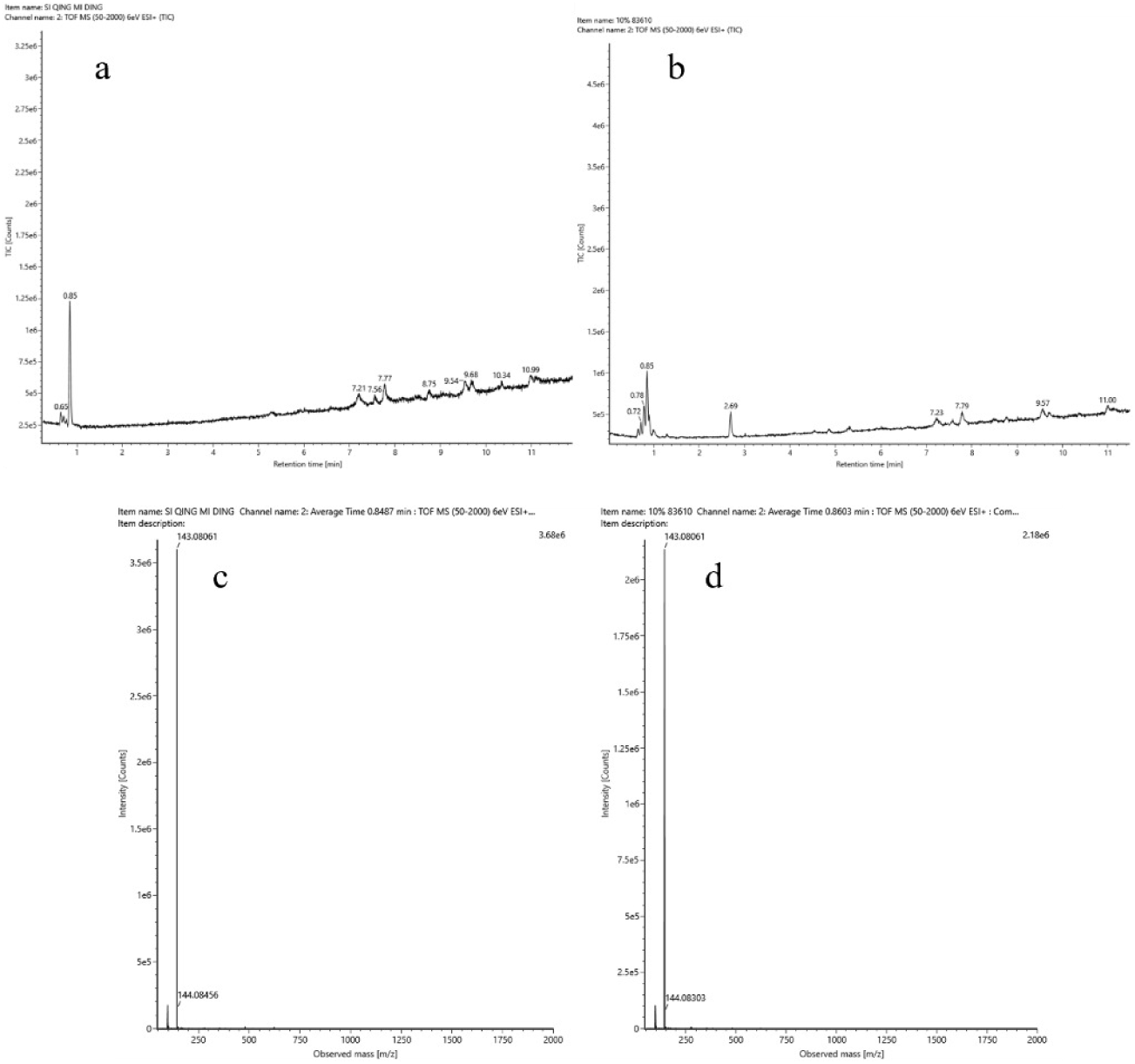
Total Ion chromatograms (TIC) and mass spectra of the sample and ectoine standard. a: Total ion chromatogram of the ectoine standard; b: Total ion chromatogram of the sample; c: Mass spectrum of the ectoine standard; d: Mass spectrum of the sample.

#### Quality Control Assessment in Targeted Metabolomics

The extracted ion chromatograms (EIC) of the standards (Fig. 10a) demonstrate satisfactory chromatographic separation, with sharp and symmetrical peaks, enabling mass spectrometric quantitative analysis of the metabolites. The relative standard deviation (RSD) of the quality control (QC) samples was less than 20% (Fig. 10b), indicating stable and reliable sample data. These experimental samples are suitable for qualitative and quantitative detection analyses of several compatible solutes.

**Fig 10.**
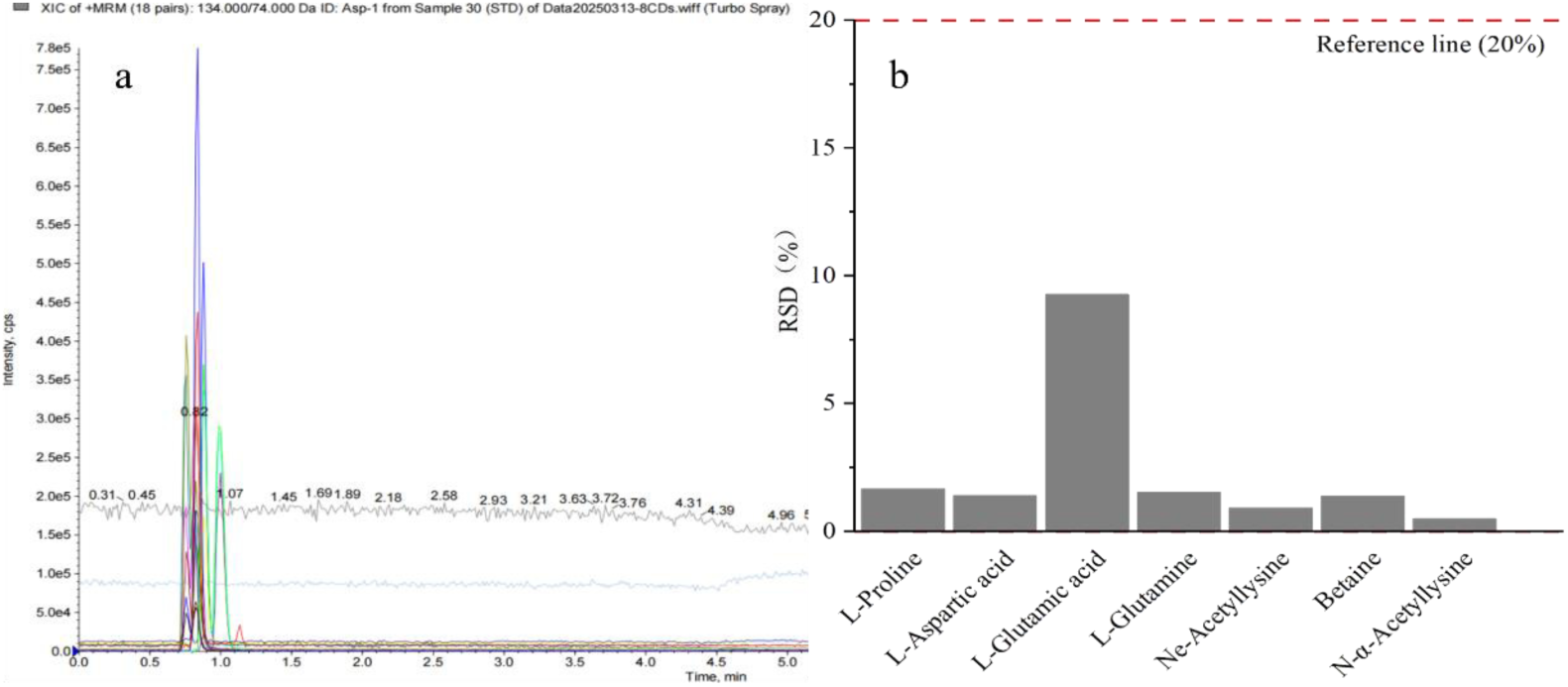
Extracted ion chromatograms and RSD plot of samples. a: Extracted ion chromatogram (EiC) of the standards; b: Relative standard deviation (RSD) of quality control (QC) samples.

#### Targeted Metabolomics Analysis

LC-MS coupled with MRM mode was employed to detect five potential compatible solutes in the samples, including Nε-Acetyl-L-lysine, and quantitative analysis of the detected compounds was performed based on standard curves of the respective reference standards (Fig. S20). As shown in Fig. 11, five compounds in the sample exhibited product ion retention times identical to those of the Nε-Acetyl-L-lysine, betaine, L-proline, L-glutamic acid, and L-glutamine standards, confirming the intracellular presence of these solutes in TRM 83610. Table 3 reveals that the levels of betaine, L-proline, L-glutamic acid, and L-glutamine gradually decreased with increasing NaCl concentrations. Notably, the Nε-Acetyl-L-lysine content increased within the 0–10% NaCl range but dropped sharply at 15% NaCl.

**Fig 11.**
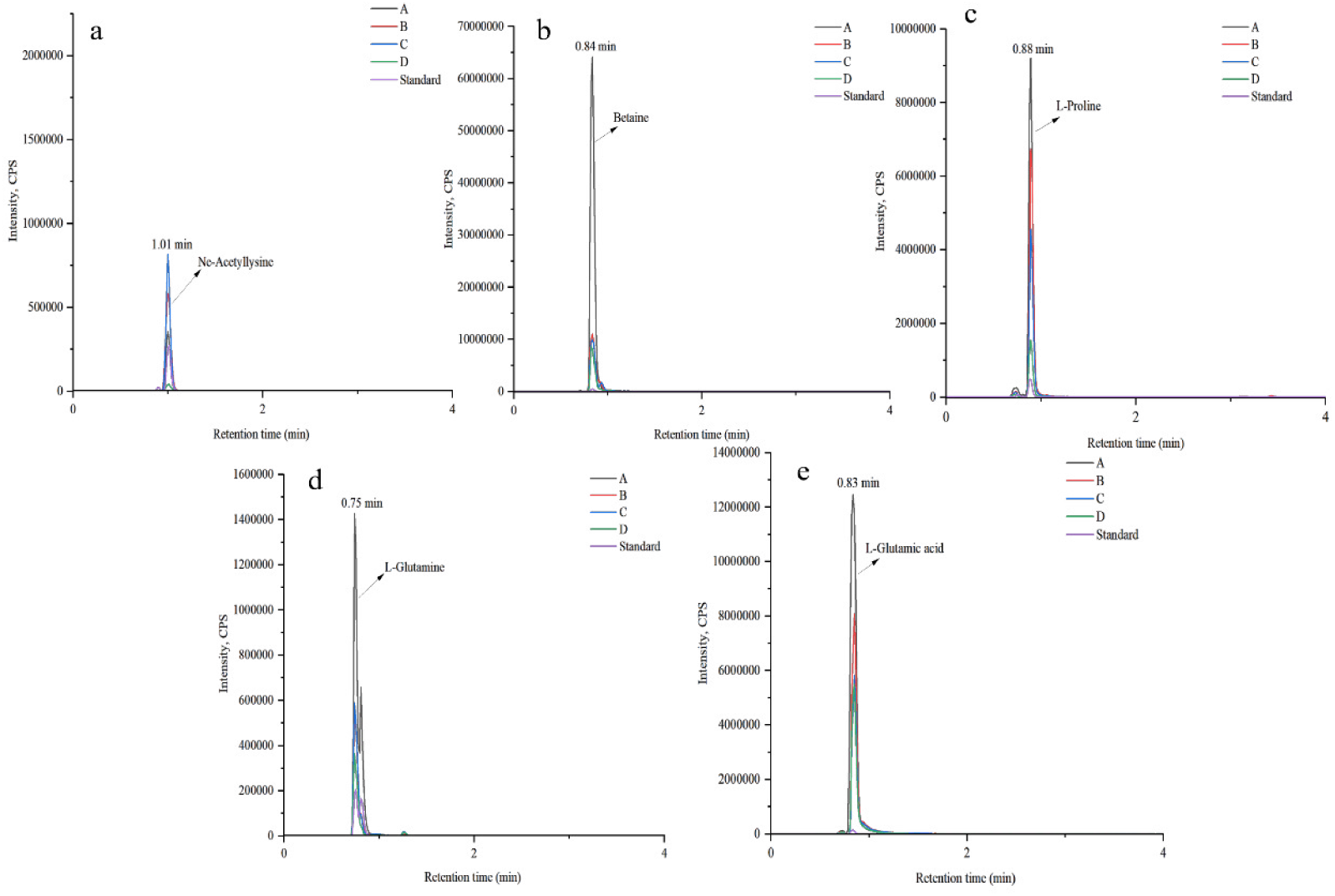
Chromatograms of the sample and five standards. a-e: Chromatograms of the standard and sample for NE-Acetyl-L-lysine, betaine, L-proline, L-glutamine, and L-glutamic acid, respectively.

**Table 3.** Content of Compatible Solutes in Sample Groups.

| ID | Name | Content (mg/kg) |  |  |  |  |
| --- | --- | --- | --- | --- | --- | --- |
|  |  | L-Proline | L-Glutamic acid | L-Glutamine | Nε-Acetyl-L-lysine | Betaine |
| 1 | A1 | 507.1 | 13890 | 77.7 | 25.4 | 5984 |
| 2 | A2 | 506.8 | 14399 | 72.9 | 28.1 | 5628 |
| 3 | A3 | 514.6 | 14302 | 48.0 | 20.5 | 5874 |
| 4 | B1 | 246.9 | 2248 | 12.2 | 27.4 | 1097 |
| 5 | B2 | 238.2 | 1700 | 8.8 | 26.4 | 999 |
| 6 | B3 | 281.7 | 1810 | 10.7 | 33.5 | 986 |
| 7 | C1 | 92.0 | 1129 | 5.77 | 42.0 | 514 |
| 8 | C2 | 147.3 | 1491 | 5.46 | 46.6 | 643 |
| 9 | C3 | 180.1 | 1835 | 8.22 | 45.4 | 818 |
| 10 | D1 | 36.9 | 999 | 2.86 | 0.63 | 389 |
| 11 | D2 | 60.3 | 1510 | 3.44 | 0.30 | 518 |
| 12 | D3 | 57.1 | 1403 | 3.68 | 0.40 | 505 |

A–D: Represent NaCl concentrations of 0, 5%, 10%, and 15%, respectively.

### Optimization of ectoine production in *B. epidermidis* using Response surface methodology

#### Plackett-Burman design results

A 12-run Plackett-Burman design was implemented using Minitab 21 software to evaluate eight factors listed in Table 4 (see Table 5 for experimental design details). Fermentation cultures were performed under the 12 distinct factor combinations specified by the design. Ectoine content in the bacterial biomass was subsequently quantified via high-performance liquid chromatography (HPLC).

**Table 4.** Factors and levels in the Plackett-Burman experimental design.

| Factor No. | Factors | Low | High |
| --- | --- | --- | --- |
| X<sub>1</sub> | Sucrose | 1 g/L | 5 g/L |
| X<sub>2</sub> | Soybean peptone | 20 g/L | 30 g/L |
| X<sub>3</sub> | Sodium glutamate | 6.76 g/L | 13.52 g/L |
| X<sub>4</sub> | Complex salt concentration | 100 g/L | 150 g/L |
| X<sub>5</sub> | Medium fill volume | 50 mL | 150 mL |
| X<sub>6</sub> | Temperature | 31°C | 37°C |
| X<sub>7</sub> | Inoculation volume | 4% | 6% |
| X<sub>8</sub> | Fermentation time | 4 d | 8 d |

**Table 5.** Plackett-Burman experimental design matrix and results.

| Serial Number | Factors |  |  |  |  |  |  |  | Production |
| --- | --- | --- | --- | --- | --- | --- | --- | --- | --- |
|  | X<sub>1</sub> | X<sub>2</sub> | X<sub>3</sub> | X<sub>4</sub> | X<sub>5</sub> | X<sub>6</sub> | X<sub>7</sub> | X<sub>8</sub> |  |
| 1 | -1 | 1 | -1 | -1 | -1 | 1 | 1 | 1 | 339.05 |
| 2 | -1 | 1 | 1 | -1 | 1 | -1 | -1 | -1 | 217.18 |
| 3 | 1 | 1 | -1 | 1 | -1 | -1 | -1 | 1 | 321.31 |
| 4 | 1 | -1 | 1 | -1 | -1 | -1 | 1 | 1 | 362.99 |
| 5 | 1 | 1 | 1 | -1 | 1 | 1 | -1 | 1 | 220.18 |
| 6 | -1 | -1 | -1 | 1 | 1 | 1 | -1 | 1 | 182.30 |
| 7 | -1 | -1 | 1 | 1 | 1 | -1 | 1 | 1 | 178.28 |
| 8 | 1 | -1 | -1 | -1 | 1 | 1 | 1 | -1 | 210.97 |
| 9 | -1 | -1 | -1 | -1 | -1 | -1 | -1 | -1 | 373.12 |
| 10 | 1 | 1 | -1 | 1 | 1 | -1 | 1 | -1 | 190.10 |
| 11 | -1 | 1 | 1 | 1 | -1 | 1 | 1 | -1 | 349.05 |
| 12 | 1 | -1 | 1 | 1 | -1 | 1 | -1 | -1 | 327.34 |

Using ectoine yield as the response variable, regression analysis was performed with Minitab 21 to establish a first-order regression equation:

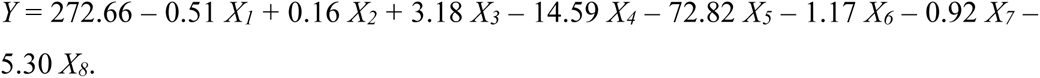

The coefficient of determination (*R*²) for the model was 98.96%, indicating excellent goodness-of-fit, and thus the equation is suitable for predicting ectoine yield. Regression analysis identified the statistical significance of the eight factors on the response variable. Among the tested factors, complex salt concentration (*P* = 0.045) and medium fill volume (*P* = 0.000) exhibited *P* < 0.05 (Table 6), confirming their statistically significant impacts on ectoine yield, with medium fill volume showing the most pronounced effect. The remaining six factors had no significant influence. The *T*-values for medium fill volume (−16.52) and complex salt concentration (−3.31) indicated negative correlations between these factors and ectoine production.

**Table 6.** Main effects analysis of Plackett-Burman design.

| Factors | Coefficient | T | P |
| --- | --- | --- | --- |
| Constant | 272.66 | 61.87 | 0.000 |
| Sucrose | -0.515 | -0.12 | 0.916 |
| Soybean peptone | 0.16 | 0.04 | 0.974 |
| Sodium glutamate | 3.18 | 0.72 | 0.523 |
| Complex salt concentration | -14.59 | -3.31 | 0.045 |
| Medium fill volume | -72.82 | -16.52 | 0.000 |
| Temperature | -1.17 | -0.27 | 0.807 |
| Inoculation volume | -0.92 | -0.21 | 0.849 |
| Fermentation time | -5.30 | -1.20 | 0.315 |

#### Steepest ascent experiment

The Plackett-Burman design revealed that complex salt concentration and medium fill volume were the critical factors influencing ectoine production in TRM83610. Both factors exhibited negative effects, necessitating appropriate reduction of their values. Step sizes were determined based on the magnitude of these factors: medium fill volume was incrementally decreased starting from 115 mL, and complex salt concentration was reduced stepwise from 125 g/L. Other conditions were set to the optimized values determined by single-factor experiments. After 6 days of fermentation, ectoine content in the biomass was quantified. Group 3 achieved the maximum ectoine yield (Table 7). Consequently, the factor levels from Group 3 (complex salt concentration: 109 g/L; medium fill volume: 85 mL) were selected as the central point for subsequent response surface optimization design.

**Table 7.** Design of the steepest ascent experiment.

| Run | Medium Fill Volume (mL) | Complex Salt Concentration (g/L) | Yield (mg/L) |
| --- | --- | --- | --- |
| 1 | 115 | 125 | 294.54 |
| 2 | 100 | 117 | 325.04 |
| 3 | 85 | 109 | 427.00 |
| 4 | 70 | 101 | 386.61 |
| 5 | 55 | 93 | 311.50 |

#### Response surface design

A central composite design (CCD) was implemented using Minitab 21 software to investigate the effects of complex salt concentration and medium fill volume on ectoine yield. Ectoine production (Y) was designated as the response variable, with complex salt concentration (X₄) and medium fill volume (X₅) as independent variables. Each factor was tested at five coded levels (−1.414, −1, 0, 1, 1.414). The factors and levels for the CCD are detailed in Table 8. Results from the CCD indicated that the fermentation conditions of 109 g/L complex salt concentration and 85 mL medium fill volume approached optimality for ectoine production (Table 9).

**Table 8.** Factors and levels of the central composite design (CCD) in response surface methodology.

| ID | Name | Level |  |  |  |  |
| --- | --- | --- | --- | --- | --- | --- |
|  |  | -1.414 | -1 | 0 | 1 | 1.414 |
| X<sub>4</sub> | Complex salt concentration (g/L) | 97.68 | 101.00 | 109.00 | 117.00 | 120.31 |
| X<sub>5</sub> | Medium fill volume (mL) | 63.79 | 70.00 | 85.00 | 100.00 | 106.21 |

**Table 9.**
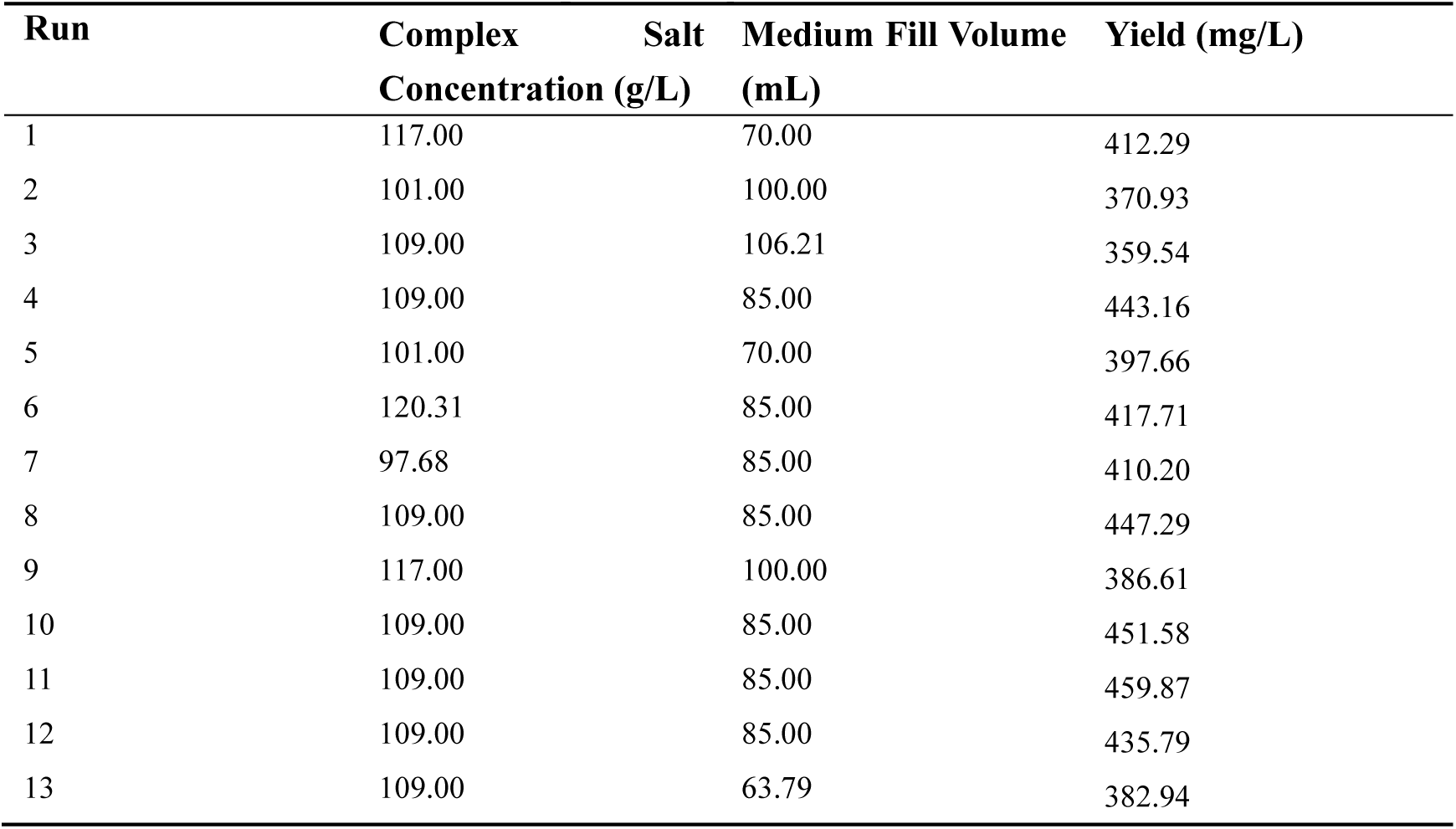
Results of the central composite design (CCD)

A quadratic polynomial regression equation was derived via binary regression fitting in Minitab 21, expressing ectoine yield (Y) as a function of complex salt concentration (X₄) and medium fill volume (X₅) in uncoded units:

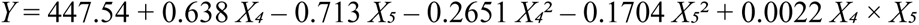

Analysis of variance (ANOVA) demonstrated a high model reliability, with an F-value of 41.87 and a P-value of 0.000.

#### Response surface analysis and validation

Response surface plots (Fig. 12) and contour plots (Fig. 13) were generated using Minitab 21 software. As shown in Fig. 11, the downward-opening response surface indicates the presence of a maximum point in the regression model. By solving the first-order partial derivatives of the regression equation, the predicted maximum ectoine yield of 448.66 mg/L was identified at 110.19 g/L complex salt concentration (X₄) and 82.92 mL medium fill volume (X₅). Balancing model predictions and practical feasibility, the optimized conditions were adjusted to 110.20 g/L complex salt concentration and 83.00 mL medium fill volume.

**Fig 12.**
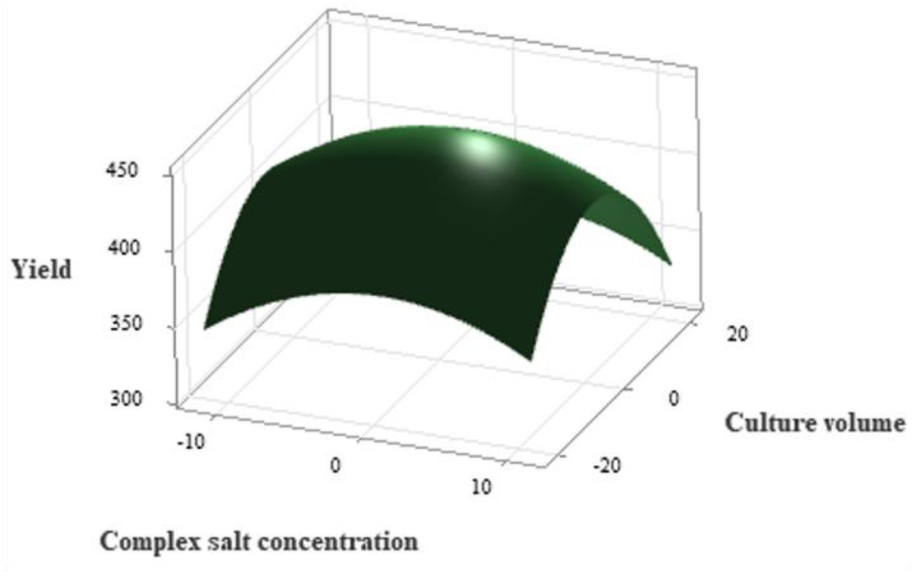
Response surface plot of complex salt concentration and medium fill volume on ectoine yield.

**Fig 13.**
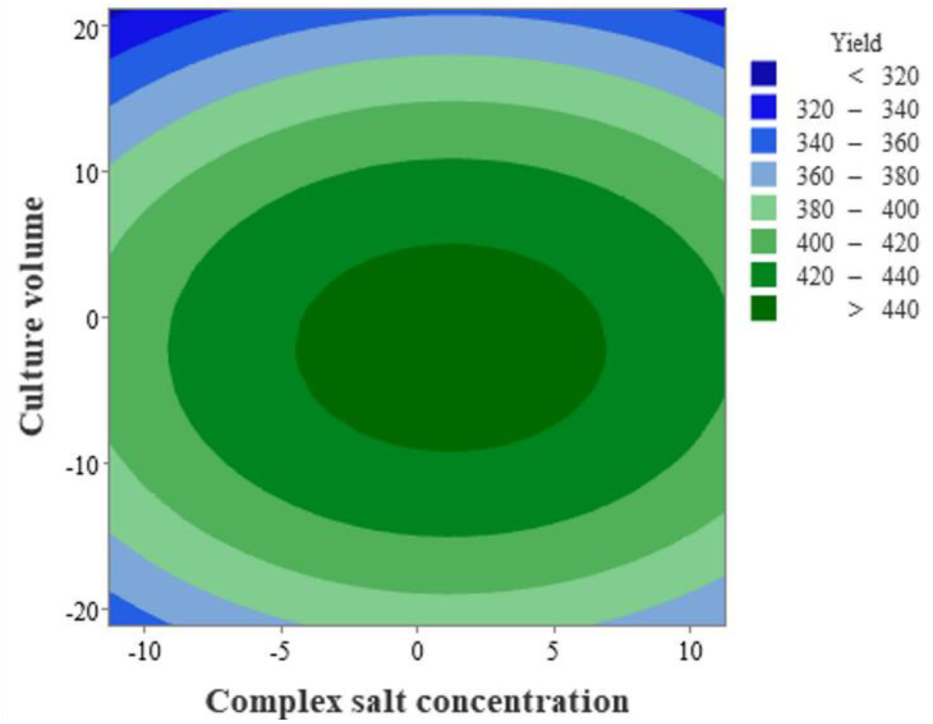
Contour plot of complex salt concentration and medium fill volume on ectoine yield.

Validation experiments were conducted using the optimized soybean peptone-sucrose medium and cultivation parameters. Ectoine content in the biomass was quantified via high-performance liquid chromatography (HPLC). The measured ectoine yield reached 440.60 mg/L, closely matching the theoretical prediction (448.66 mg/L) and representing a 6.22-fold increase over the initial yield of 70.75 mg/L.

## DISCUSSION

Metabolomic analysis identified six compatible solutes, predominantly ectoine, whose concentrations varied significantly with NaCl levels, albeit with distinct trends. Previous studies indicate that halotolerant microorganisms synthesize diverse compatible solutes depending on environmental conditions^[46]^, dynamically adjusting metabolic pathways to regulate solute concentrations and maintain osmotic balance. A proposed “two-phase” salt tolerance strategy ^[47, 48]^ suggests an initial rapid response phase, where cells accumulate readily available solutes (e.g., betaine, L-glutamic acid) to counteract osmotic stress and enable proliferation^[49, 50]^, followed by synthesis of more complex solutes like ectoine and Nε-acetyl-L-lysine for long-term protein stabilization. Aston et al. ^[51]^ observed salinity-dependent shifts in compatible solute profiles in *Halomonas campisalis*. We hypothesize that *Brevibacterium epidermidis* TRM83610 initially accumulates betaine, L-glutamic acid, L-glutamine, and L-proline as temporary osmoprotectants during early NaCl stress, later transitioning to ectoine as the primary solute. Notably, aspartate, glutamate, and glutamine directly participate in ectoine biosynthesis, while betaine and proline may indirectly support its synthesis. Their declining concentrations, inversely correlated with ectoine accumulation, suggest metabolic reprogramming under salt stress ^[52, 53]^.

Correlation analysis revealed strong negative associations between ectoine and 14 organic acids/derivatives. While amino acid consumption has been linked to ectoine synthesis ^[54–56]^, this study implicates non-amino organic acids in this process. Intriguingly, Nε-acetyl-L-lysine showed no significant correlation with other solutes and decreased sharply at 15% NaCl, suggesting roles beyond osmoprotection, potentially in protein acetylation to enhance stress resistance^[57, 58]^.

*B. epidermidis* TRM83610 employs multifaceted NaCl adaptation strategies. Elevated 5-hydroxyindoleacetate (a serotonin oxidation product) under high salinity indicates oxidative stress mitigation ^[59, 60]^, while upregulated antimicrobial compounds suggest enhanced ecological competitiveness ^[61, 62]^. Metabolic pathway enrichment (e.g., amino acid biosynthesis, D-amino acid metabolism) highlights coordinated regulation of ectoine synthesis and stress-responsive pathways. Antimicrobial/antioxidant metabolites, though distributed across pathways, predominantly localized to “Metabolic pathways” and “Biosynthesis of plant secondary metabolites,” underscoring the strain’s integrated osmotic, oxidative, and competitive adaptation mechanisms.

This study provides a preliminary evaluation of the application potential of *B. epidermidis* TRM83610. Firstly, the strain exhibits strong ectoine synthesis capability. Wild-type strains typically demonstrate low ectoine yields, generally not exceeding 270 mg/L ^[63, 64]^. For instance, Hong et al. ^[65]^ isolated two halophilic strains from Jilantai Salt Lake soil with maximum ectoine yields of 80.35 mg/L and 97.89 mg/L, respectively. Yao et al.^[66]^ reported an ectoine yield of 92.41 mg/L for *Halomonas ventosae* Al12T AY268080. Tian et al.^[67]^ increased the ectoine yield of the wild-type strain *Halomonas campaniensis* to 456.82 mg/L through fermentation optimization.

Wang et al. ^[68]^obtained a *Halomonas campaniensis* XH26 mutant via UV-induced mutagenesis, achieving ectoine yields ranging from 380 to 660 mg/L. In this study, fermentation optimization of the wild-type *B. epidermidis* TRM 83610 resulted in a significantly higher ectoine yield of 440.62 mg/L, representing a 6.22-fold increase over the initial yield of 70.75 mg/L, demonstrating its robust ectoine production capacity. Furthermore, untargeted metabolomics annotated multiple antimicrobial active substances, a survival strategy advantageous for becoming the dominant strain in open fermentation processes^[62, 69]^. Notably, 5-Hydroxyectoine, often a major byproduct of ectoine biosynthesis ^[70]^, was not detected in this study, thereby reducing downstream recovery costs and complexity.

Secondly, *B. epidermidis*TRM 83610 shows promise for broader applications. Nε-Acetyl-L-lysine has been proposed for biotechnology applications ^[71]^. The enrichment of plant growth-promoting metabolites suggests its potential as a Plant Growth-Promoting Bacterium (PGPB). Additionally, the possible production of various antimicrobial and anti-inflammatory active substances further expands its application prospects. Finally, it is noteworthy that Azetidomonamide A was annotated as a different metabolite (DM) via machine learning. To our knowledge, this compound has previously only been reported in *Pseudomonas aeruginosa*^[72]^. In this study, the relative abundance of Azetidomonamide A initially increased and subsequently decreased with rising NaCl concentrations. Previous research has demonstrated that Azetidomonamide A participates in regulating biofilm formation and pigment synthesis in Pseudomonas aeruginosa, and its biosynthesis is modulated by quorum sensing (QS) ^[73]^. Therefore, we hypothesize that *B. epidermidis* TRM 83610 may perceive and respond to environmental changes through a QS mechanism to modulate physiological activities, warranting further investigation.

### Conclusions

We identified six compatible solutes in *B. epidermidis* TRM83610, including first-reported Nε-acetyl-L-lysine. The strain employs multipronged NaCl adaptation: ectoine-dominated osmotic regulation, antioxidant synthesis, and antimicrobial production. Its high ectoine yield (440.62 mg/L) and metabolic versatility position it as a promising platform for microbial cell factories, with applications spanning biomanufacturing, agriculture, and biomedicine.

## MATERIALS AND METHODS

### Strain and Cultivation

*Brevibacterium epidermidis* TRM83610, isolated from Mangya Jade Lake (Qinghai-Tibet Plateau), was preserved at the Tarim Basin Biological Resource Conservation and Utilization Key Laboratory.

Fermentation medium: 10 g yeast extract, 7.5 g acid-hydrolyzed casein peptone, NaCl (as required), 1 L distilled water.

### Untargeted metabolomics

Fermentation media were separately prepared with NaCl concentrations of 0, 5%, 10%, and 15%, each concentration comprising six biological replicates. Following sterilization via autoclaving at 121°C for 20 minutes and subsequent cooling, media were inoculated with 2% (v/v) seed culture. Cultivation proceeded for 6 days at 37°C with 150 rpm orbital shaking. Bacterial cells were then harvested by centrifugation at 4000 rpm, washed twice with isotonic NaCl solution through repeated centrifugation cycles, and the final pellet was stored overnight at −80°C. Lyophilized biomass was pulverized into homogeneous powder. Untreated *B. epidermidis*TRM 83610 (0 NaCl) served as the control group, while samples exposed to 5%, 10%, and 15% NaCl constituted treatment groups.

Untargeted metabolomic analysis was conducted by Shanghai Personal Biotechnology Co., Ltd. (Personalbio). Aliquots of 50 mg sample powder were transferred to 2 mL centrifuge tubes containing 1 mL of pre-chilled 50% methanol. After vortex mixing for 30 s, samples underwent flash-freezing in liquid nitrogen for 5 minutes followed by thawing at room temperature. Homogenization was performed using a high-throughput tissue grinder at 55 Hz for 60 s, with 2-3 repetition cycles. Subsequent centrifugation at 12,000 rpm (4°C, 15 min) yielded supernatants, 800 μL of which were vacuum-concentrated to complete dryness. The residue was reconstituted in 150 μL of 50% methanol containing 5 ppm 2-chloro-L-phenylalanine as internal standard, vortexed for 30 s, and centrifuged (12,000 rpm, 4°C, 10 min). Final supernatants were filtered through 0.22 μm membranes into vials. Quality control (QC) samples were prepared by pooling 10-20 μL aliquots from each sample to monitor instrumental stability and data reliability.

Chromatographic separation employed an ACQUITY UPLC HSS T3 column (100Å, 1.8 μm, 2.1 × 100 mm) maintained at 40°C with 0.4 mL/min flow rate and 2 μL injection volume. Mobile phases consisted of (A) 0.1% formic acid in water and (B) acetonitrile containing 0.1% formic acid. The gradient program was: 0-1 min (5% A, 95% B); 1-7 min (5-95% A); 7-8 min (95% A, 5% B); 8.1-12 min (5% A, 95% B). High-resolution mass spectrometry operated in data-dependent acquisition (DDA) mode under Xcalibur software control (v4.7, Thermo Scientific) using a HESI ion source with spray voltage set at 3.5 kV. Key parameters included: sheath gas 40 arb, auxiliary gas 15 arb, capillary temperature 325°C, auxiliary gas heater 300°C. Full-scan MS1 spectra (m/z 100-1000) were acquired at 60,000 resolution (AGC target standard, max IT 100 ms), with top-4 precursors selected for MS2 fragmentation at 15,000 resolution using 30% normalized collision energy, dynamic exclusion of 8 s, and automatic maximum injection time.

### Targeted metabolomics

Qualitative detection of ectoine was performed by the Analytical Testing Center of Tarim University. LC-MS conditions were as follows: An ACQUITY UPLC-BEH C18 column (1.7 μm, 2.1 × 100 mm) was employed for ultrahigh-performance liquid chromatography. Key parameters included capillary voltage 2.50 kV, source temperature 100°C, desolvation temperature 500°C, cone gas flow 50 L/h, desolvation gas flow 800 L/h, with mass scanning range set at m/z 50-2000 and scan time 0.20 s. Targeted metabolomic analysis was conducted by Yanxuan Biotechnology (Hangzhou) Co., Ltd. Samples were subsampled from those used in untargeted metabolomics analysis, with three biological replicates per group. LC-MS analysis based on selective multiple reaction monitoring (MRM) technology utilized a Shimadzu Nexera X2 LC-30AD UHPLC system. Mobile phases consisted of (A) 0.1% formic acid aqueous solution and (B) acetonitrile containing 0.1% formic acid. Chromatographic separation proceeded at 40°C column temperature with 300 μL/min flow rate and 1 μL injection volume. Mass spectrometric detection was performed on a 5500 QTRAP mass spectrometer (AB Sciex) in positive ion mode. ESI source parameters were configured as: source temperature 550°C; ion source gas 1 (GS1): 55 psi; ion source gas 2 (GS2): 55 psi; curtain gas (CUR): 35 psi; ion spray voltage (IS): 5500 V. Detection was carried out in MRM mode.

### Experimental design

Single-factor tests evaluated 10 variables: carbon/nitrogen sources, sodium glutamate (0.02–0.10 mol/L), salt (25–175 g/L), fermentation time (1–9 d), temperature (25– 43°C), agitation (100–200 rpm), pH (5.5–10.5), inoculum size (1–8%), and medium fill volume (50–250 mL).

Plackett-Burman and steepest ascent designs identified critical factors (complex salt concentration, fill volume) for response surface optimization (Minitab 21).

### Data Analysis

Raw data were preprocessed and subjected to quality control using the XCMS package in R, filtering out metabolites with RSD > 30%. Metabolite identification was performed by matching against public databases (HMDB, MassBank, LipidMaps, mzCloud, KEGG) and Biocode’s in-house metabolite library, with identification confidence levels set at Level 2 or above.

Relative abundance analysis of metabolites was conducted using ggplot2 (v3.4). Pairwise comparative differential analysis of sample data was performed using the Ropls R package. Multi-group comparative differential analysis employed the PMCMRplus (v1.9), Pheatmap (v1.0), and clusterProfiler (v4.6) packages in R. Multi-group differential volcano plots and association network diagrams were generated on the Biocode GeneCloud platform (https://www.genescloud.cn, accessed April 1, 2025).

Standard curves, chromatograms of samples versus five compatible solute standards, DMs, and bar charts for fermentation optimization were plotted using Origin 2024 software. The metabolic network diagram was created with iodraw (https://www.iodraw.com/). Response surface plots and contour plots were generated using Minitab 21. Data are expressed as mean values.

## ACKNOWLEDGMENTS

This work was supported by the Third Xinjiang Scientific Expedition Program (2022xjkk150307), OpenFunding Project of State Key Laboratory of Microbial Metabolism (MMLKF22-01).

The authors extend their gratitude to Scientific Compass (www.shiyanjia.com) for providing invaluable assistance with the targeted metabolomic analysis.

## DATA AVAILABILITY

The datasets generated from untargeted metabolomics and targeted metabolomics analyses in this study are available via the Metabolights database under accession numbers MTBLS12624 and MTBLS12638, respectively.

